# Rapid identification of *Biomphalaria* spp. and diagnosis of *Schistosoma mansoni* infestation using MALDI-TOF mass spectrometry

**DOI:** 10.1101/2025.06.16.659835

**Authors:** Diara Sy, Lionel Almeras, Adama Zan Diarra, Souleymane Doucoure, Yacine Mbere Sarr, Coralie L’Ollivier, Pape Mouhamed Gaye, Pape Ibrahima Ndiaye, Bruno Senghor, Doudou Sow, Cheikh Sokhna, Stephane Ranque

**Affiliations:** Aix-Marseille University, IRD, AP-HM, SSA, RITMES, 13005, Marseille, France; EMR MINES Infectious, Neglected, and Emerging Diseases in the South, Aix-Marseille University, Research Institute for Development, 13005 Marseille, France; EMR MINES: Infectious, Neglected, and Emerging Diseases in the South, Research Institute for Development, International Campus of the Institute of Research for Development, Université Cheikh Anta Diop de Hann, Dakar BP 1386, Senegal; IHU-Méditerranée Infection, 13005, Marseille, France; Parasitology and Entomology Unit, Department of Microbiology and Infectious Diseases, Armed Forces Biomedical Research Institute, 13005 Marseille, France; Department of Parasitology-Mycology, Health Sciences Training and Research Unit, Université Gaston Berger, Saint Louis BP 234, Senegal

**Keywords:** Schistosomiasis, *Biomphalaria*, *S. mansoni*, MALDI-TOF MS

## Abstract

This study explores the use of Matrix-Assisted Laser Desorption/Ionization Time-of-Flight mass spectrometry (MALDI-TOF MS) to identify and differentiate *Biomphalaria* snails infected with the parasite *S. mansoni*, which causes schistosomiasis. The study was conducted on two snail species, *Biomphalaria pfeifferi* (collected in the field in Senegal) and *Biomphalaria glabrata* (a laboratory strain). The snails were infected in the laboratory with *S. mansoni* miracidia, and their infection was confirmed by cercariae emission tests and quantitative PCR. MALDI-TOF MS was then used to analyse proteins from infected and uninfected snails to identify spectral differences. Based on protein profiles, the results of MALDI-TOF mass spectrometry made it possible to accurately differentiate between S. mansoni-infected snails and uninfected snails. An increase in the number of peaks detected and their intensity was observed for the spectra of S. mansoni-infected snails compared to uninfected snails. The application of principal component analysis to these mass spectrometry profiles confirmed the discrimination between the two groups according to their infection status. In addition, specific discriminating peaks were identified for each snail species, allowing for the distinction of infected from uninfected snails. The present study revealed, for the first time, that MALDI-TOF MS appears to be a rapid, reliable, and specific tool for the diagnosis of schistosomiasis in snails, offering promising prospects for the surveillance and control of this disease in endemic areas. However, further work is needed to establish a MALDI-TOF MS reference spectra database specific to *Schistosoma* parasites and to standardise sample collection, storage, and preparation in order to apply this technique in the field.

## Introduction

Schistosomiasis is a parasitic infection caused by contact with surface water contaminated by the parasite of the genus *Schistosoma* (De Boni *et al*., 2021; Riaz *et al*., 2023). This parasitic disease is very serious and affects more than 190 million people worldwide (Shi *et al*., 2022). Seventy-eight countries are currently facing problems resulting from human schistosomiasis, the majority of them in sub-Saharan Africa (Tchuem Tchuenté *et al*., 2017). Freshwater snails can be intermediate hosts for many parasites of medical and veterinary interest (Dreyfuss and Rondelaud, 2011), and the genus *Biomphalaria* contributes to the transmission of *Schistosoma mansoni*, responsible for intestinal schistosomiasis (Chu, 1976). *Biomphalaria pfeifferi* is a freshwater snail which is widely distributed in sub-Saharan Africa and Madagascar, as well as in some areas of the Sahara and Southwest Asia (Brown, 1994). Another species, *Biomphalaria glabrata* is native to South America (Dejong *et al*., 2003). Meanwhile, in Senegal, the only species of the genus *Biomphalaria* is *B. pfeifferi*, an intermediate host of the parasite *S. mansoni* (Moyroud *et al*., 1982). It is, therefore, clear that several strategies are needed to control schistosomiasis, including controlling the infestation dynamics of molluscs in endemic contact areas (King, 2009).

Several diagnostic methods for parasite surveillance have been developed to monitor the evolutionary dynamics of schistosomiasis in endemic areas (McPhail *et al*., 2022). These include manual cercariometry (Sato *et al*., 2018; Theron, 1979), positive phototropism (Klock, 1961), the superposition technique (Sandt, 1972), and centrifugation (Butler *et al*., 1967). However, to detect infestation with these methods, the parasite must be in its final stage of evolution. Techniques such as qPCR (Jothikumar *et al*., 2015) and molecular detection and identification (Pennance *et al*., 2020) have emerged to detect parasites at an early stage. These two methods, however, cannot be implemented in the field and require significant preparation time. Recently, an innovative approach has emerged for the identification and diagnosis of pathogens and their vectors, based on protein profiling: matrix-assisted laser desorption/ionization time-of-flight mass spectrometry (MALDI-TOF MS) (Costa *et al*., 2024; Sevestre *et al*., 2021). MALDI-TOF MS has proven to be a rapid, reliable, and specific method for identifying parasites, bacteria, viruses, and arthropod vectors, representing significant potential in the field of clinical microbiology and disease control (Sánchez-Juanes *et al*., 2022). However, it has also enabled us to identify the intermediate host snails of the parasite *Schistosoma* (Hamlili *et al*., 2021). In addition, Huguenin *et al*. were able to identify and differentiate different species of parasites of the genus *Schistosoma* (Huguenin *et al*., 2023). The primary objective of this study was to discriminate between the two species of *Biomphalaria, B. pfeifferi* and *B. glabrata*, using MALDI-TOF MS and then to assess whether MALDI-TOF MS could detect the *S. mansoni* parasite in the snails.

## Materials and methods

### Snail collection

A field mission to collect *B. pfeifferi* snails, which are likely to transmit the *S. mansoni* parasite, was conducted in 2022 in the Saint Louis region of northern Senegal, an area known for perennial transmission of schistosomiasis. The samples were sent to the IRD Hann Mariste laboratory in Dakar, where they were identified (Brown, 1994) and tested for cercariae emission to separate infested snails, specifically those that emitted cercariae. The isolated cercariae were used to infest hamsters in groups of 200 cercariae per hamster to increase the likelihood of testing positive for schistosomiasis after 90 days. Snails that tested negative for cercariae emission after 45 days in the laboratory were reared in our facility until the F1 generation was obtained, which is well adapted to laboratory conditions and free of *S. mansoni*. The *B. glabrata* snail used in our experiment was a Brazilian strain from the Naval Medical Research Institute (NMRI) and was kindly donated by the IHPE laboratory.

### Ethics statement

The experimental protocol was approved by the CNERS Senegal Ethics Committee (Reference No. 00000019) and received clearance from the head of Nagoya in Senegal (Reference No. 001339). The protocol complies with national and international guidelines on animal welfare.

### Laboratory infestation

Laboratory infestation is a common procedure for maintaining the *S. mansoni* parasite cycle. Cercariae collected from *S. mansoni*-positive *B. pfeifferi* in the field were used to infect three laboratory-bred hamsters through the cutaneous route. After three months (90 days), one of the three hamsters was sacrificed to collect *S. mansoni* eggs trapped in visceral tissues and adult worms in the vena cava. The other two hamsters were kept for future infestations. To achieve this, the eggs were placed in distilled water and exposed to natural light for up to 30 minutes. The adult worms were preserved for further study.

### *Biomphalaria* infestation in the laboratory

After egg hatching, *S. mansoni* miracidia were observed under a ×100 magnifying glass and collected using a Pasteur pipette. The F1 generation of *B. pfeifferi* snails, measuring between 3 mm and 5 mm, and the *B. glabrata* snails were then infected with these *S. mansoni* miracidia.

To do so, we used a 24-well pillbox for each species, placing one individual and five miracidia in the shade for three hours. Finally, the individuals were collected in tanks and then placed in semi-natural conditions (with day-night alternation). According to the literature, cercariae emission tests can begin on the 23^rd^ day after infection (Capron *et al*., 1965) and can continue until the 30^th^ day. Individuals shedding cercariae were considered positive for infestation during this period. In this study, cercariae shedding tests were assessed between day 21 and day 30 of infestation.

### PCR-based *S. mansoni* detection

The presence of the *S. mansoni* parasite in snails was confirmed by the qPCR system as previously described (Wichmann *et al*., 2009). Briefly, we used the forward primers SRA1 and antisense primers SRS2 and the probe SRP. The PCR mixing conditions were 20 μl of reaction mix containing 5 μl of template DNA, 3.5 μl of sterile ultrapure water, 0.5 μl of each primer, 0.5 μl of probe, and 10 μl of Master Mix (product ref). Amplification was performed on a CFX96 thermal cycler (BIO-RAD, Hercules, CA, USA). The PCR programme was initiated by a three-minute purification at 95 °C, followed by ten minutes’ denaturation at 95 °C, and 40 cycles of 30 seconds of denaturation and hybridisation at 55 °C for 30 seconds, maintained at 4 °C (Wichmann *et al*., 2009). In each run, a negative control (sterile water) and a positive control (DNA extracted from *S. mansoni* worms) were used. PCR results were considered positive if the cycle threshold (CT) was less than 35 cycles.

### MALDI TOF MS-based *S. mansoni* detection

#### Snail preparation

The soft part from the shell was carefully removed and the foot was dissected under a Leica ES2 10x/30x stereomicroscope using a sterile slide. The remaining parts of the snail were stored at -20 °C for further study. The dissected part was first rinsed with 70% ethanol and then distilled water for two minutes. It was dried, deposited on a sterile filter paper, and cut into small pieces (Hamlili *et al*., 2021). The foot samples were then added to an extraction solution composed of a mixture of 70% formic acid (Sigma-Aldrich, Lyon, France), 50% acetonitrile (Fluka, Buchs, Switzerland), and 80% HPLC water. For each sample, 30 μL of the mixed solution and glass beads (Sigma-Aldrich, St. Louis, Missouri, USA) was homogenised using a Tissue Lyser II (Qiagen, Germany) with optimised parameters (three one-minute cycles at a frequency of 30 Hertz), as described previously (Yssouf *et al*., 2013). All homogenates were centrifuged at 2000 g for 30 seconds, and 1.5 μL of each supernatant was spotted onto a steel target plate (Bruker Daltonics GmBh, Bremen, Germany) in four spots. One microlitre of a CHCA matrix suspension composed of saturated alpha-cyano-4-hydroxycinnamic acid (Sigma, Lyon, France), 50% acetonitrile, 2.5% trifluoroacetic acid (Aldrich, Dorset, UK), and 47.5% of HPLC grade water was directly deposited onto each spot on the target plate to allow co-crystallisation, then dried at room temperature before being inserted into the MALDI-TOF MS instrument (Bruker Daltonics) (Nebbak *et al*., 2016).

#### Analysis of MALDI-TOF MS spectra

Mass spectrometry analysis was performed using a Microflex LT MALDI-TOF mass spectrometer (Bruker Daltonics) with the Flex Control software (Bruker Daltonics). Measurements were performed in linear positive ion mode in a 2 kDA–20 kDa mass range. Each spectrum corresponds to ions obtained from 240 laser shots performed in six regions of a single spot. Spectral profiles were visualised using Flex Analysis, version 3.3, and then exported to the MALDI Biotyper (Bruker Daltonics) software version 3.0 and ClinProTools (Bruker Daltonics) for further data processing (smoothing, baseline subtraction, peak selection). The reproducibility of MS spectra was assessed by comparing the Main Spectrum Profile (MSP) obtained from the four spots of each sample with the MALDI Biotyper (Bruker Daltonics). The composite correlation index (CCI) tool of the MALDI Biotyper was also used to assess spectral variations within and between snails which were infected with *S. mansoni* and those that were not (Diarra *et al*., 2019). In addition, ClinProTools was used to identify discriminatory peaks between infected and non-infected snails not only within species but also between the two species. The most discriminatory common peaks between infected and non-infected snails were analysed with ClinProTools to compare the two groups and estimate the impact of infestation. The ClinProTools default parameters for spectrum preparation were applied as described by Briolant *et al*. (Briolant *et al*., 2020). Based on the peak list obtained per species, the five and eight most intense peaks of snails infected by *B. pfeifferi* and *B. glabrata*, respectively were selected for inclusion in the genetic algorithm (GA) model. The peaks selected by the operator provided a recognition capacity (RC) value related to the highest cross-validation (CV) value. The presence or absence of all discriminatory peak masses generated by the GA model was checked by comparing the average spectra of each species per body part.

#### DNA sequence-based snail species confirmation

To confirm the identification of *Biomphalaria* species, we sequenced three infected and three uninfected specimens of each species that displayed a good MALDI-TOF MS spectrum and a high MALDI Biotyper (Bruker Daltonics) log score value (LSV) ≥ 2. To identify the snail species, we sequenced the following gene fragments: COX (CO1 (LC1490): GGTCAACAAATCATAAAGATATTGG; CO2 (HCO2198): TAAACTTCAGGGTGACCAAAAAATCA); ITS (ETTS1: TGCTTAAGTTCAGCGGGT; ETTS10: GCATACTGCTTTGAACATCG) (Kane *et al*., 2008), and 18S (18S F: AGTATGGTTGCAAAGCTGAAACTTA; 18S R: TACAAAGGGCAGGGACGTAAT) (Webster *et al*., 2006), as previously described. To confirm the amplification of DNA, the PCR products were run on a 1% agarose gel at 180 V, 400 mA for 15 minutes. For a PCR to be considered positive, the detection of a 363-base pair band was required. Positive samples were purified and sequenced using the same primers with BigDye version 1-1 Cycle Sequencing Ready Reaction Mix (Applied Biosystems, Foster City, CA) and run on an ABI 3100 automated sequencer (Applied Biosystems). Sequences were assembled and corrected using the ChromasPro (version 1.34) software (Technelysium Pty. Ltd., Tewantin, Australia) and identified using BLAST queries (https://blast.ncbi.nlm.nih.gov/Blast.cgi) against the NCBI GenBank nucleotide database.

## Results

### Field sampling and laboratory infestation of hamsters and *Biomphalaria*

A total of 15 snails identified as *B. pfeifferi* were collected in the field in the Saint Louis region. At the Dakar laboratory, nine of the 15 *B. pfeifferi* (60%) were found to be dead. Of the six remaining live snails, three (50%) tested positive for cercarial emission in the laboratory. These three positive snails were exposed to sunlight to recover as many cercariae as possible for hamster infestation. To maintain the parasite cycle, three hamsters were infected with *S. mansoni* cercariae. One of the hamsters was sacrificed, and the *S. mansoni* miracidia hatching from the eggs were used to infect 48 snails of the genus *Biomphalaria* (24 *B. pfeifferi* and 24 *B. glabrata*). After a 30-day rearing period in the laboratory, all 48 snails were exposed to natural light for 15 minutes. A total of 40 out of 48 (83.3%) snails (20 *B. pfeifferi* and 20 *B. glabrata*) emitted cercariae and were therefore considered to be infected with *S. mansoni*. All cercariae-positive snails were confirmed using the *S. mansoni*-specific qPCR, with CT values ranging from 10 to 21.

#### Snail species DNA sequence-based confirmation

Seven snails specimens, five of which were infected with *S. mansoni*, and three uninfected snails of each species, were subjected to standard PCR and sequencing using three different genes (Cox, ITS and 18S). BLAST analysis of *B. pfeifferi* and *B. glabrata* sequences showed coverage percentages of 98% to 100% and identity percentages ranging from 97% to 100% with the sequences of their counterparts in GenBank (Table 1).

**Table 1.**
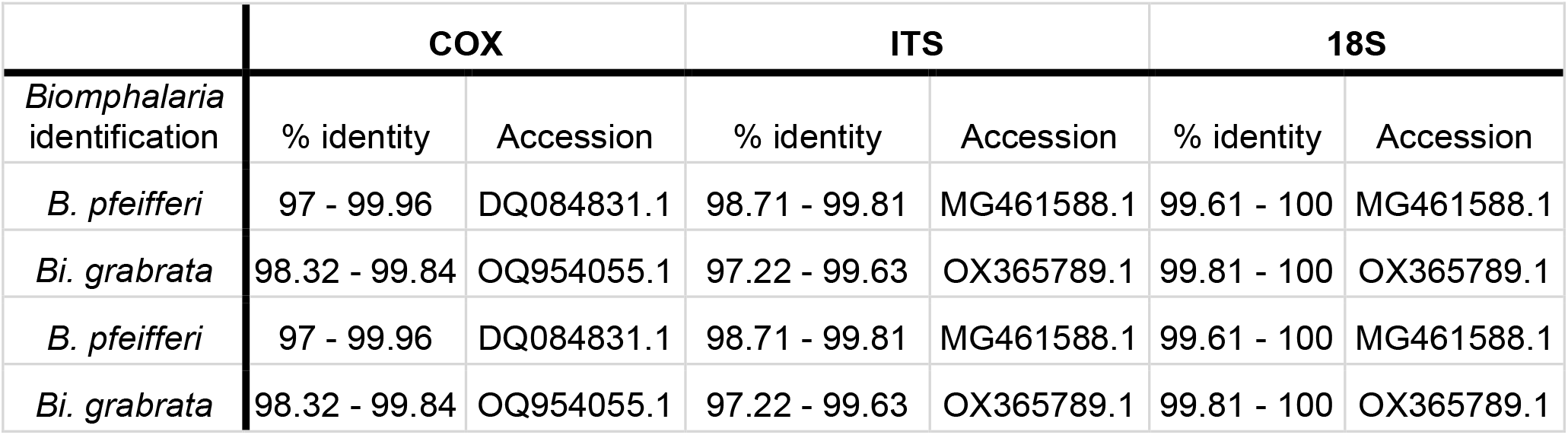
Molecular identification of *Biomphalaria* snails.

|  | COX |  | ITS |  | 18S |  |
| --- | --- | --- | --- | --- | --- | --- |
| *Biomphalaria*<br>identification | % identity | Accession | % identity | Accession | % identity | Accession |
| *B. pfeifferi* | 97 - 99.96 | DQ084831.1 | 98.71 - 99.81 | MG461588.1 | 99.61 - 100 | MG461588.1 |
| *Bi. grabrata* | 98.32 - 99.84 | OQ954055.1 | 97.22 - 99.63 | OX365789.1 | 99.81 - 100 | OX365789.1 |
| *B. pfeifferi* | 97 - 99.96 | DQ084831.1 | 98.71 - 99.81 | MG461588.1 | 99.61 - 100 | MG461588.1 |
| *Bi. grabrata* | 98.32 - 99.84 | OQ954055.1 | 97.22 - 99.63 | OX365789.1 | 99.81 - 100 | OX365789.1 |

### MALDI-TOF MS spectra analysis

The MALDI-TOF MS spectra of uninfected *Biomphalaria* snails were selected to assess the ability of this tool to discriminate between the two species. We obtained high intensity spectra, i.e., greater than 2000 au, when 20 uninfected *Biomphalaria* snails were subjected to MALDI TOF MS. These spectra appeared visually reproducible for each species (Figure 2A). To confirm the reproducibility of the spectra, cluster analyses were performed, resulting in the MSP dendrogram (Figure 2B). Three specimens of each species were selected. The clustering of specimens of the same species on the same branch with a high distance between them and the absence of intertwining species demonstrate the reproducibility and specificity of each *Biomphalaria* species protein profile.

**Figure 1:**
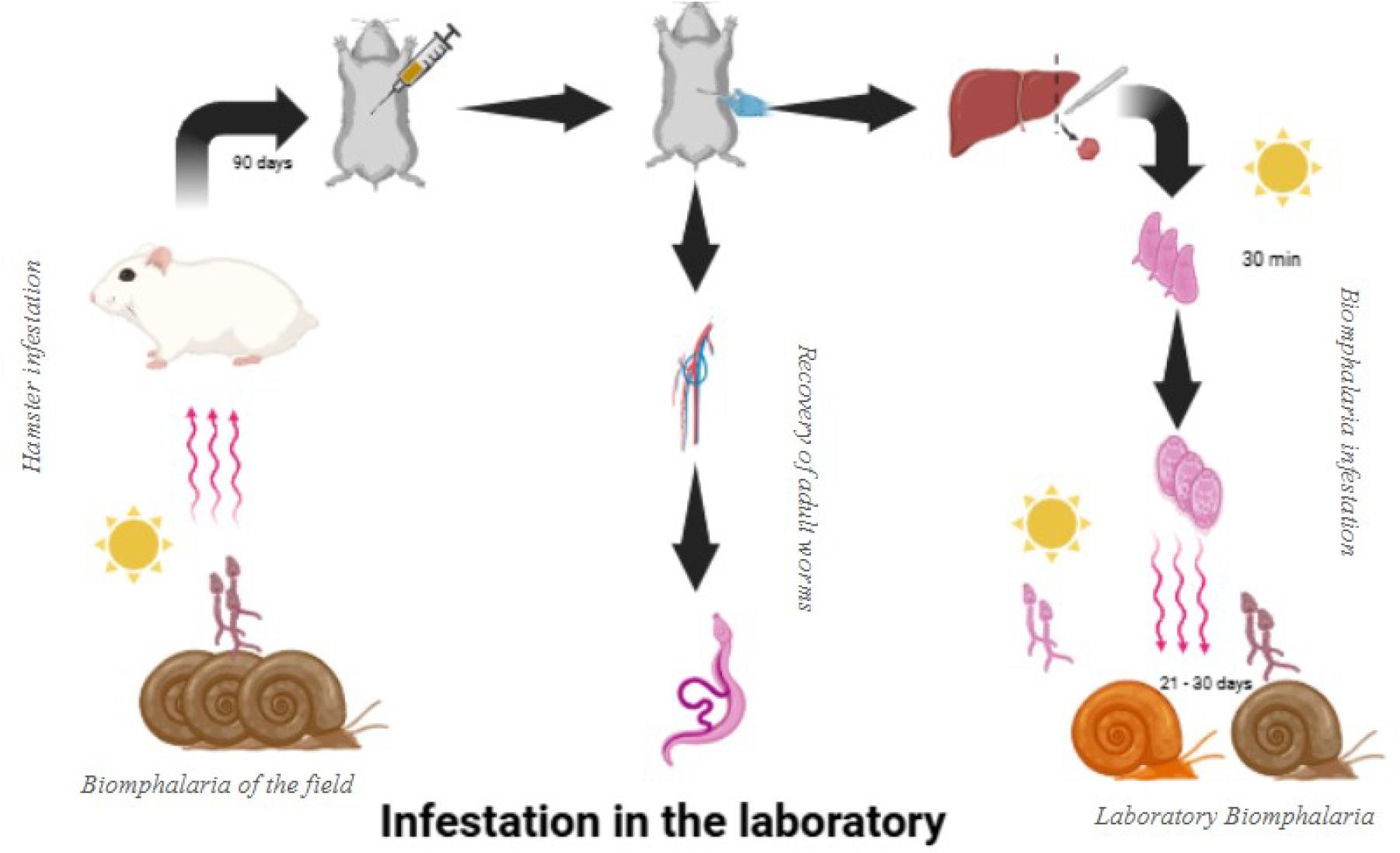
Infestation of snails of the genus *Biomphalaria* in the laboratory

**Figure 2.**
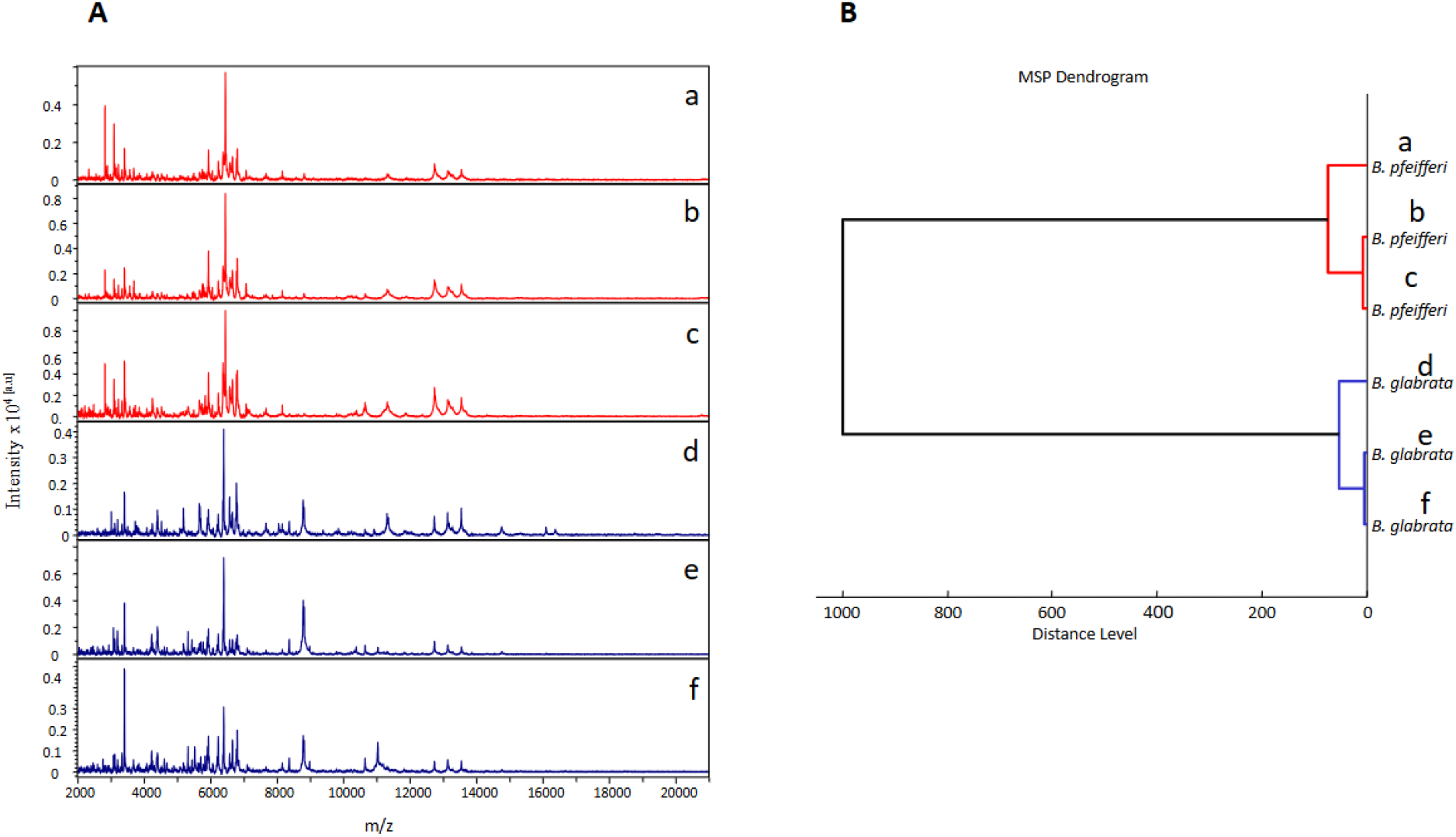
Identification of *Biomphalaria* by MALDI-TOF MS profiling. (A) Comparison of MALDI-TOF MS spectra obtained from the snails stipes of three distinct specimens of *B. pfeifferi* (a, b, c) and *B. glabrata* (d, e, f). (m/z: mass-to-charge ratio). (B) MSP dendrogram of the MALDI-TOF MS spectra of three specimens per species. The dendrogram was created using the Biotyper v3.0 software, and the distance units correspond to the relative similarity of the MS spectra.

Since the correct classification of the snail species relies primarily on the intensity of the resulting MS spectra, the five most intense peaks were selected from the uninfected *Biomphalaria* snails. This resulted in a total of eight peaks discriminating between the *Biomphalaria* species (Table 2). These eight peaks were subjected to the genetic algorithm (GA) model of ClinProTools 2.2 software. The combination of the presence or absence of these five most intense peaks displayed Cross-Validation (CV) and Recognition Capacity (RC) values equal to 100%. These five peaks were also evaluated against the infested groups. This enabled us to determine the percentage of compatibility between the groups. Values of 84% and 80% were obtained for infested *B. glabrata* and infested *B. pfeifferi*, respectively.

**Table 2.** List of the top eight mass peaks per *Biomphalaria* species using feet as biological material.

| Average peak intensity (a.u.)* |  |  |  |
| --- | --- | --- | --- |
| MS peak number (§) | m/z (Da) | *B. glabrata* | *B. pfeifferi* |
| 1 | 2814.36 | 2.72 | **18.58** |
| 2 | 3400.21 | **22.46** | **14.63** |
| 3 | 5925.07 | **10.1** | **16.12** |
| 4 | 6221.43 | **10.33** | 6.96 |
| 5 | 6366.49 | 8.46 | **13.39** |
| 6 | 6387.48 | **19.09** | 11.67 |
| 7 | 6438.18 | 3.35 | **33.66** |
| 8 | 6794.58 | **10.51** | 12.51 |

§List of MS peaks used to distinguish *Biomphalaria* species, based on analysis by the ClinProTools genetic algorithm model. \*The top five mass peaks per *Biomphalaria* species that discriminate between the two species are shown in bold. (Da: Daltons; m/z: mass/charge; a.u.: arbitrary unit.)

The two *Biomphalaria* species formed two distinct MALDI-TO spectra groups. In Figure 3, the infected group is in green and the uninfected group is in red. The separation of infected and uninfected snails was much clearer in the *B. glabrata* species, with ∼85% of variance being explained, while only ∼80% of the variance was explained in the *B. pfeifferi* species.

**Figure 3:**
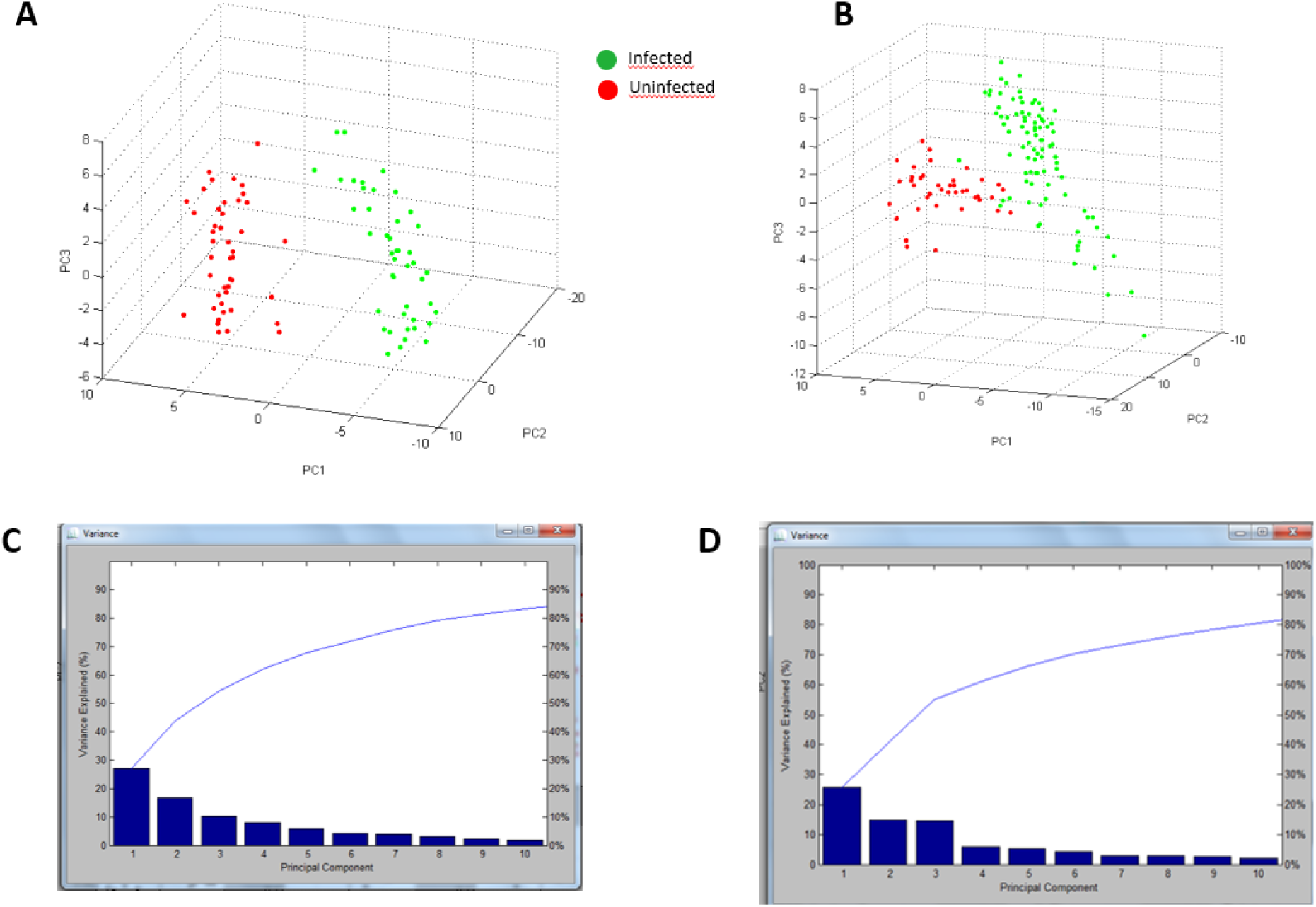
Comparison of the different spectra obtained from the two Biomphalaria species, infected or not infected by *S. mansoni*, using principal component analysis (PCA). A. *B. glabrata* infected (n=20, green dots) or not infected (n=10, red dots) with S. mansoni. B. *B. pfeifferi* infected (n=20, green dots) or not infected (n=10, red dots) with *S. mansoni*. C. Expression curve of variance versus principal component of *B. glabrata*. C. Expression curve of variance versus principal component of *B. pfeifferi*.

Within the *B. glabrata* snail group, 16 peaks differentiated snails infected with the *S. mansoni* parasite from uninfected snails. Statistical analyses showed a *P-*value less than .005 (*P* < .005) in the PTTA Analysis of Variance test. This indicates that these peaks are informative and discriminatory on the infestation parameter. Using ClinProTools software, principal component analysis generated a cross-validation percentage of 97.67% and a recognition ability of 100%.

**Table 3.** The 16 most discriminating MALDI-TOF MS peaks, selected with ClinProTools software, discriminating infected *B. glabrata* snails from uninfected ones.

| Average peak intensity (a.u.)* |  |  |  |  |  |
| --- | --- | --- | --- | --- | --- |
| MS peak number§ | m/z (Da) | PTTA | *B. glabrata* neg (BgN) | *B. glabrata* pos (BgP) | Ratio BgP/BgN |
| 1 | 2072.63 | < 0.000001 | 2.12 | 5.36 | 2.53 |
| 2 | 2217.21 | < 0.000001 | 2.42 | 5.41 | 2.24 |
| 3 | 2304.48 | < 0.000001 | 2.22 | 5.33 | 2.40 |
| 4 | 2399.19 | < 0.000001 | 2.49 | 9.82 | 3.94 |
| 5 | 3628.45 | < 0.000001 | 1.41 | 15.79 | 11.20 |
| 6 | 3644.45 | < 0.000001 | 1.63 | 6.33 | 3.88 |
| 7 | 4802.11 | < 0.000001 | 1.02 | 35.53 | 34.83 |
| 8 | 4818.03 | < 0.000001 | 1.11 | 10.15 | 9.14 |
| 9 | 5175.48 | < 0.000001 | 5.13 | 2.29 | 0.45 |
| 10 | 5311.79 | < 0.000001 | 3.42 | 1.28 | 0.37 |
| 11 | 6387.3 | < 0.000001 | 19.09 | 6.13 | 0.32 |
| 12 | 8355.94 | < 0.000001 | 5.17 | 2.13 | 0.41 |
| 13 | 8779.91 | < 0.000001 | 8.2 | 3.35 | 0.41 |
| 14 | 8804.39 | < 0.000001 | 7.11 | 2.81 | 0.40 |
| 15 | 10640.59 | < 0.000001 | 2.54 | 0.86 | 0.34 |
| 16 | 13129.7 | < 0.000001 | 5.72 | 2.37 | 0.41 |

§List of MS peaks used to distinguish *Biomphalaria glabrata* species, based on analysis by the ClinProTools genetic algorithm model. \* The 17 main mass peaks per species of *Biomphalaria glabrata*, which allow to differentiate between the two groups of infected/non-infected snails, Da: Daltons; m/z: mass/charge; a.u.: arbitrary unit, PTTA Analysis of Variance test *P*-value

**Table 4.** The 17 most discriminating MALDI-TOF MS peaks, selected with ClinProTools software, between infected *B. pfeifferi* snails and uninfected ones.

| Average peak intensity (a.u.)* |  |  |  |  |  |
| --- | --- | --- | --- | --- | --- |
| MS peak number§ | m/z (Da) | PTTA | *B. pfeifferi* neg (BPn) | *B. pfeifferi* pos (BPp) | Ratio BPp/BPn |
| 1 | 4568.01 | 0.00231 | 1.69 | 5.97 | 0.28 |
| 2 | 4802.27 | 0.00834 | 1.12 | 5.28 | 0.21 |
| 3 | 4872.64 | 0.00000136 | 1.26 | 3.95 | 0.32 |
| 4 | 4889.05 | 0.0000166 | 1.98 | 4.26 | 0.46 |
| 5 | 4927.1 | 0.00000557 | 1.24 | 5.03 | 0.25 |
| 6 | 5487.01 | < 0.000001 | 3.79 | 1.61 | 2.35 |
| 7 | 5765.23 | < 0.000001 | 4.49 | 2.02 | 2.22 |
| 8 | 5829.13 | < 0.000001 | 4.82 | 1.38 | 3.49 |
| 9 | 5901.72 | < 0.000001 | 4.76 | 2.37 | 2.01 |
| 10 | 5924.64 | < 0.000001 | 16.12 | 6.72 | 2.40 |
| 11 | 6127.44 | < 0.000001 | 2.27 | 4.65 | 0.49 |
| 12 | 6437.76 | < 0.000001 | 33.66 | 8.28 | 4.07 |
| 13 | 6457.5 | < 0.000001 | 9.13 | 4.46 | 2.05 |
| 14 | 6583.76 | < 0.000001 | 7.26 | 2.99 | 2.43 |
| 15 | 7060.81 | < 0.000001 | 5.26 | 1.93 | 2.73 |
| 16 | 10641.45 | 0.00000529 | 1.83 | 0.87 | 2.10 |
| 17 | 13130.64 | < 0.000001 | 4.2 | 1.67 | 2.51 |

§List of MS peaks used to distinguish between *Biomphalaria glabrata* species, based on analysis by the ClinProTools genetic algorithm model. \* The 17 main mass peaks per species of *Biomphalaria glabrata*, which allow to distinguish between the two infected/non-infected groups, Da: Daltons; m/z: mass/charge; a.u.: arbitrary unit, PTTA Analysis of Variance test *P*-value

## Discussion

First, MALDI TOF MS made it possible to discriminate between two snail species of the same genus. Specific and reproducible spectra were obtained for each species with 100% reliability. In addition, we identified *B. glabrata* for the first time, using MALDI TOF MS with good quality spectra (LSV > 2). This confirms the efficiency and reliability of MALDI-TOF in identifying snail species, as confirmed by Hamlili *et al*., (Hamlili *et al*., 2021) in their previous study.

Second, we were able to clearly and accurately discriminate between infected and uninfected *Biomphalaria* snails using MALDI-TOF MS. MALDI-TOF, initially limited to identifying intermediate host snails (Hamlili *et al*., 2021), is capable of differentiating parasite infestations within the host. Since this device is a tool based on the ionisation of an organism’s proteins (Diarra *et al*., 2019), MALDI-TOF detected the presence of other proteins on samples that were positive for cercariae emission and real-time PCR, which is different from the proteins detected in uninfected snails. This resulted in a difference in spectra between the group of infected and uninfected snails. Thus, the visualised spectra confirm a difference in protein expression between infected *Biomphalaria* and uninfected *Biomphalaria*. The reproducibility in the spectra of infected snails indicates the presence of the same proteins in these snails. The same observations were made in the uninfected snails. Given that there is no MALDI-TOF spectra database to identify *Schistosoma* parasites, the parasite species remain to be identified.

The validation percentages and intra-species recognition capacity obtained confirm that MALDI-TOF is indeed capable of differentiating *Schistosoma* infestation within the same species. On the one hand, we observed different peaks, which indicate the expression of distinct proteins, and on the other we observed similar peaks in both infected and uninfected groups. In this case, the distinction lies in the intensity and regularity of expression in all individuals. In other words, the presence of the parasite can modulate the expression of certain proteins in the snail, thus making it possible to differentiate between infected snails and those that are not. Similarly, it has been demonstrated that MALDI-TOF MS could differentiate filaria-infected mosquitoes from uninfected mosquitoes, with a specificity of 100% and a sensitivity of 92% (Tahir *et al*., 2017).

The slight difference in the validation percentages and the recognition capacity between *B. pfeifferi* and *B. glabrata* may be explained by the origin of the collected samples. The *B. pfeifferi* samples were collected in the field, while our *B. glabrata* strain had been raised in the laboratory for years. Thus, uncontrolled environmental conditions such as the physicochemical parameters of the water and the instability of the climate, but also water pollution can influence MALDI-TOF MS spectra and contribute to the differences between the spectra of field and laboratory snails. Indeed, MALDI-TOF MS has been found to be capable of tracing the geospatial origin of snails (Gaye *et al*., 2023).

## Conclusion

This study demonstrates the effectiveness of Matrix-Assisted Laser Desorption/Ionization Time-of-Flight (MALDI-TOF) mass spectrometry as an innovative tool for the identification and differentiation between snails of the genus *Biomphalaria* infected with *Schistosoma mansoni*, the causative agent of schistosomiasis. By combining speed, reliability, and specificity, this method offers promising prospects for diagnosing and monitoring this parasitic disease in endemic areas, particularly in sub-Saharan Africa, where schistosomiasis is a major public health problem. The good quality MALDI-TOF spectra generated in this study will be used to enrich our in-house MALDI-TOF spectral database and further blind tests are planned to assess the capacity of MALDI-TOF MS to diagnose *Schistosoma* parasite infestations of snails collected in the field and to propose a standardised sample collection, storage, and preparation protocol. Finally, this technological advance will contribute towards developing an integrated approach aimed at reducing the public health impact of schistosomiasis and contributing to global efforts to eradicate this disease by 2030, in line with WHO targets.

## Authors’ contribution

DS (Diara Sy), SD, and DS (Doudou Sow) conceived and designed the study. DS (Diara Sy), BS, and YMS participated in data collection. DS (Diara Sy), LC and PMG carried out laboratory activities. DS (Diara Sy), AZD, LA and PIN performed data analyses and wrote the first draft of the manuscript. AZD, CS, SR, LA, SD, and DS (Doudou Sow) revised the article.

## Funding

This work was supported by the University Hospital Institute (IHU) Mediterranean Infection, 13005, Marseille, France; EMR MINES: Infectious, Neglected, and Emerging Diseases in the South, Campus International Institute of Research for Development, Université Cheikh Anta Diop de Hann, Dakar BP 1386, Senegal.

## Availability of data and materials

All relevant data generated during this study are included in the manuscript.

## Notes

### Competing Interest Statement

The authors have declared no competing interest.

## Bibliography

Briolant, S., Costa, M.M., Nguyen, C., Dusfour, I., Pommier de Santi, V., Girod, R., Almeras, L., 2020. Identification of French Guiana anopheline mosquitoes by MALDI-TOF MS profiling using protein signatures from two body parts. PLoS One 15, e0234098. 10.1371/journal.pone.0234098

Brown, D.S., 1994. Freshwater Snails Of Africa And Their Medical Importance, 0 ed. CRC Press. 10.1201/9781482295184

Butler, J.M., Ferguson, F.F., Palmer, J.R., 1967. New field device for quantitative recovery of Schistosoma mansoni cercariae. Public Health Rep (1896).82, 250–252. 10.2307/4592983

Capron, A., Deblock, S., Biguet, J., Clay, A., Adenis, L., Vernes, A., n.d. 1965. Contribution a 1’etude experimentale de la bilharziose a Schistosoma haematobium. Bull. Wld. Hlth Org 32, 755–778. https://iris.who.int/handle/10665/267228

Chu, K.Y., 1976. The validity of baseline data for measuring incidence rates of Schistosoma haematobium infection in the molluscicided area, UAR-0049 project. Annals of Tropical Medicine & Parasitology 70, 365–367. 10.1080/00034983.1976.11687133

Costa, M.M., Corbel, V., Ben Hamouda, R., Almeras, L., 2024. MALDI-TOF MS Profiling and Its Contribution to Mosquito-Borne Diseases: A Systematic Review. Insects 15, 651. 10.3390/insects15090651

De Boni, L., Msimang, V., De Voux, A., Frean, J., 2021. Trends in the prevalence of microscopically-confirmed schistosomiasis in the South African public health sector, 2011– 2018. PLoS Negl Trop Dis 15, e0009669. 10.1371/journal.pntd.0009669

Dejong, R.J., Morgan, J.A.T., Wilson, W.D., Al‐Jaser, M.H., Appleton, C.C., Coulibaly, G., D’Andrea, P.S., Doenhoff, M.J., Haas, W., Idris, M.A., Magalhães, L.A., Moné, H., Mouahid, G., Mubila, L., Pointier, J. ‐P., Webster, J.P., Zanotti‐Magalhães, E.M., Paraense, W.L., Mkoji, G.M., Loker, E.S., 2003. Phylogeography of Biomphalaria glabrata and B. pfeifferi, important intermediate hosts of Schistosoma mansoni in the New and Old World tropics. Molecular Ecology 12, 3041–3056. 10.1046/j.1365-294X.2003.01977.x

Diarra, A.Z., Laroche, M., Berger, F., Parola, P., 2019. Use of MALDI-TOF MS for the Identification of Chad Mosquitoes and the Origin of Their Blood Meal. Am J Trop Med Hyg 100, 47–53. 10.4269/ajtmh.18-0657

Dreyfuss, G., Rondelaud, D., 2011. Les mollusques dans la transmission des helminthoses humaines et vétérinaires. 10.4267/2042/48064

Gaye, P.M., Doucouré, S., Sow, D., Sokhna, C., Ranque, S., 2023. Identification of Bulinus forskalii as a potential intermediate host of Schistosoma hæmatobium in Senegal. PLoS Negl Trop Dis 17, e0010584. 10.1371/journal.pntd.0010584

Hamlili, F.Z., Thiam, F., Laroche, M., Diarra, A.Z., Doucouré, S., Gaye, P.M., Fall, C.B., Faye, B., Sokhna, C., Sow, D., Parola, P., 2021. MALDI-TOF mass spectrometry for the identification of freshwater snails from Senegal, including intermediate hosts of schistosomes. PLoS Negl Trop Dis 15, e0009725. 10.1371/journal.pntd.0009725

Huguenin, A., Kincaid-Smith, J., Depaquit, J., Boissier, J., Ferté, H., 2023. MALDI-TOF: A new tool for the identification of Schistosoma cercariae and detection of hybrids. PLoS Negl Trop Dis 17, e0010577. 10.1371/journal.pntd.0010577

Jothikumar, N., Mull, B.J., Brant, S.V., Loker, E.S., Collinson, J., Secor, W.E., Hill, V.R., 2015. Real-Time PCR and Sequencing Assays for Rapid Detection and Identification of Avian Schistosomes in Environmental Samples. Appl Environ Microbiol 81, 4207–4215. 10.1128/AEM.00750-15

Kane, R.A., Stothard, J.R., Emery, A.M., Rollinson, D., 2008. Molecular characterization of freshwater snails in the genus Bulinus: a role for barcodes? Parasit Vectors 1, 15. 10.1186/1756-3305-1-15

King, C.H., 2009. Toward the Elimination of Schistosomiasis. New England Journal of Medicine 360, 106–109. 10.1056/NEJMp0808041

Klock, J.W., 1961. A method for the direct quantitative recovery of Schistosoma mansoni cercariae from natural waters of Puerto Rico. Bull World Health Organ 25, 738–740. https://iris.who.int/handle/10665/267536

McPhail, B.A., Froelich, K., Reimink, R.L., Hanington, P.C., 2022. Simplifying Schistosome Surveillance: Using Molecular Cercariometry to Detect and Quantify Cercariae in Water. Pathogens 11, 565. 10.3390/pathogens11050565

Moyroud, J., Breuil, J., Dulat, Ch., Coulanges P., 1982. Les mollusques, hôtes intermediaires desbilharzioses humaines a madagascar. Etat actuel desconnaissances. Archives-Institut-Pasteur-de-Madagascar 50, 39–65. https://www.pasteur.mg/wp-content/uploads/2015/08/Archives-Institut-Pasteur-de-Madagascar-1982-50-39-65.pdf

Nebbak, A., Willcox, A.C., Bitam, I., Raoult, D., Parola, P., Almeras, L., 2016. Standardization of sample homogenization for mosquito identification using an innovative proteomic tool based on protein profiling. Proteomics 16, 3148–3160. 10.1002/pmic.201600287

Pennance, T., Archer, J., Lugli, E.B., Rostron, P., Llanwarne, F., Ali, S.M., Amour, A.K., Suleiman, K.R., Li, S., Rollinson, D., Cable, J., Knopp, S., Allan, F., Ame, S.M., Webster, B.L., 2020. Development of a Molecular Snail Xenomonitoring Assay to Detect Schistosoma haematobium and Schistosoma bovis Infections in their Bulinus Snail Hosts. Molecules 25, 4011. 10.3390/molecules25174011

Riaz, S., Ahmed, H., Kiani, S.A., Afzal, M.S., Simsek, S., Celik, F., Wasif, S., Bangash, N., Naqvi, S.K., Zhang, J., Cao, J., 2023. Knowledge, attitudes and practices related to neglected tropical diseases (schistosomiasis and fascioliasis) of public health importance: A cross-sectional study. Front Vet Sci 10, 1088981. 10.3389/fvets.2023.1088981

Sánchez-Juanes, F., Calvo Sánchez, N., Belhassen García, M., Vieira Lista, C., Román, R.M., Álamo Sanz, R., Muro Álvarez, A., Muñoz Bellido, J.L., 2022. Applications of MALDI-TOF Mass Spectrometry to the Identification of Parasites and Arthropod Vectors of Human Diseases. Microorganisms 10, 2300. 10.3390/microorganisms10112300

Sandt, D.G., 1972. Evaluation of an overlay technique for the recovery of Schistosoma mansoni cercariae. Bull World Health Organ 47, 125–127. https://iris.who.int/handle/10665/263496

Sato, M.O., Rafalimanantsoa, A., Ramarokoto, C., Rahetilahy, A.M., Ravoniarimbinina, P., Kawai, S., Minamoto, T., Sato, M., Kirinoki, M., Rasolofo, V., De Calan, M., Chigusa, Y., 2018. Usefulness of environmental DNA for detecting Schistosoma mansoni occurrence sites in Madagascar. Int J Infect Dis 76, 130–136. 10.1016/j.ijid.2018.08.018

Sevestre, J., Diarra, A.Z., Laroche, M., Almeras, L., Parola, P., 2021. Matrix-assisted laser desorption/ionization time-of-flight mass spectrometry: an emerging tool for studying the vectors of human infectious diseases. Future Microbiol 16, 323–340. 10.2217/fmb-2020-0145

Shi, L., Zhang, J.-F., Li, W., Yang, K., 2022. Development of New Technologies for Risk Identification of Schistosomiasis Transmission in China. Pathogens 11, 224. 10.3390/pathogens11020224

Tahir, D., Almeras, L., Varloud, M., Raoult, D., Davoust, B., & Parola, P. 2017. Assessment of MALDI-TOF mass spectrometry for filariae detection in Aedes aegypti mosquitoes. PLoS Neglected Tropical Diseases, 11(12), e0006093. 10.1371/journal.pntd.0006093

Tchuem Tchuenté, L.-A., Rollinson, D., Stothard, J.R., Molyneux, D., 2017. Moving from control to elimination of schistosomiasis in sub-Saharan Africa: time to change and adapt strategies. Infect Dis Poverty 6, 42. 10.1186/s40249-017-0256-8

Theron, A., 1979. A differential filtration technique for the measurement of schistosome cercarial densities in standing waters. Bull World Health Organ 57, 971–975. https://iris.who.int/handle/10665/261951

Webster, B., Southgate, V., Littlewood, D., 2006. A revision of the interrelationships of Schistosoma including the recently described Schistosoma guineensis☆. International Journal for Parasitology 36, 947–955. 10.1016/j.ijpara.2006.03.005

Wichmann, D., Panning, M., Quack, T., Kramme, S., Burchard, G.-D., Grevelding, C., Drosten, C., 2009. Diagnosing Schistosomiasis by Detection of Cell-Free Parasite DNA in Human Plasma. PLoS Negl Trop Dis 3, e422. 10.1371/journal.pntd.0000422

Yssouf, A., Flaudrops, C., Drali, R., Kernif, T., Socolovschi, C., Berenger, J.-M., Raoult, D., Parola, P., 2013. Matrix-Assisted Laser Desorption Ionization–Time of Flight Mass Spectrometry for Rapid Identification of Tick Vectors. J Clin Microbiol 51, 522–528. 10.1128/JCM.02665-12

